# Golgi-localized exo-β1,3-galactosidases involved in AGP modification and root cell expansion in Arabidopsis

**DOI:** 10.1101/2020.02.13.947820

**Authors:** Pieter Nibbering, Bent L. Petersen, Mohammed Saddik Motawia, Bodil Jørgensen, Peter Ulvskov, Totte Niittylä

**Affiliations:** Department of Forest Genetics and Plant Physiology, Umeå Plant Science Centre, Swedish University of Agricultural Sciences, 901 83 UMEÅ; Department of Plant and Environmental Sciences, University of Copenhagen, DK-1871 Frederiksberg C, Denmark

## Abstract

Plant arabinogalactan proteins (AGPs) are a diverse group of cell surface- and wall-associated glycoproteins. Functionally important AGP glycans are synthesized in the Golgi apparatus, but the relationships between their glycosylation, processing, and functionality are poorly understood. Here we report the identification and functional characterization of two Golgi-localized exo-β-1,3-galactosidases from the glycosyl hydrolase 43 (GH43) family in *Arabidopsis thaliana*. *GH43* loss of function mutants exhibit root cell expansion defects in sugar-containing growth media. This root phenotype is associated with an increase in the extent of AGP cell wall association, as demonstrated by Yariv phenylglycoside dye quantification and comprehensive microarray polymer profiling of sequentially extracted cell walls. Recombinant GH43 characterization showed that the exo-β-1,3-galactosidase activity of GH43s is hindered by β-1,6 branches on β-1,3-galactans. In line with this steric hindrance, the recombinant GH43s did not release galactose from cell wall extracted glycoproteins or AGP rich gum arabic. These results show that Arabidopsis GH43s are involved in AGP glycan biosynthesis in the Golgi, and suggest their exo-β-1,3-galactosidase activity influences AGP and cell wall matrix interactions, thereby adjusting cell wall extensibility.

## Introduction

Plant cell growth is dictated by turgor-driven extension of the primary cell wall. Oriented cellulose biosynthesis and localised wall deposition create dynamic cell wall mechanics that enable turgor-driven anisotropic cell growth (Cosgrove, 2005). Studies on the molecular interactions in the primary cell wall matrix have led to models in which covalent and non-covalent interactions between matrix components facilitate anisotropic cell expansion (Bashline et al., 2014). The primary cell wall polysaccharide matrix contains cellulose, hemicellulose and pectin as well as enzymes and structural proteins (Ellis et al., 2010; Harholt et al., 2010; Rose and Lee, 2010; Scheller and Ulvskov, 2010). The cell wall-associated glycoproteins include arabinogalactan proteins (AGPs), which are proposed to have diverse roles during cell expansion, including as structural components, modifiers of the extracellular matrix, and ligands for cell surface receptors (Showalter et al., 2010; Xue et al., 2017).

Glycans account for 90-98% of the molecular weight of AGPs, and are thought to be critical for their functionality (Showalter, 1993). Characterisation of the enzymes responsible for AGP glycosylation can elucidate AGP functions and functional redundancies (Showalter et al., 2010; Showalter and Basu, 2016). AGP glycosylation is initiated by proline hydroxylation in the ER and is then advanced in the Golgi apparatus by specialized glycosyl transferases (GTs) (Rose and Lee, 2010; Hijazi et al., 2014). The glycosyl transferase family 31 (GT31) is responsible for the synthesis of the hydroxyproline galactose and the β-1,3-galactan backbone of AGPs (Basu et al., 2015; Basu et al., 2015; Ogawa-Ohnishi and Matsubayashi, 2015; Suzuki et al., 2017). Defects in β-1,3-galactan biosynthesis are associated with various developmental phenotypes including defects in cell expansion (Basu et al., 2015; Ogawa-Ohnishi and Matsubayashi, 2015; Suzuki et al., 2017). The side chains, which branch off from the β-1,3-galactan back bone via β-1,6 linkages, are synthesized by GALT29A and/or GT31 family enzymes (Geshi et al., 2013; Dilokpimol et al., 2014). Further side chain residues are attached by other GTs: GT77 enzymes add α-1,3/1,5-arabinose units (Gille et al., 2013), by GT14 attaches β-1,6 glucuronic acid units (Knoch et al., 2013; Dilokpimol et al., 2014), and GT37 adds α-1,2-fucose units (Wu et al., 2010; Tryfona et al., 2014). These GTs acting on AGPs are expected to generate a glycan structure consisting of a β-1,3 galactan backbone with β-1,6 galactan branches containing diverse other sugars.

Despite the advances in identifying GTs involved in AGP glycosylation, the *in vivo* structural diversity of mature arabinogalactans is huge and largely uncharacterised. The glycan structure is also likely to vary between AGPs, tissues, and even cell types, as indicated by differences in extractability, expression during specific growth stages, and the expression of the glycosylation machinery (Pennell et al., 1991; Showalter et al., 2010). Nuclear magnetic resonance (NMR) analysis of the glycans attached to synthetic AGP motif repeats expressed in tobacco Bright Yellow-2 cells revealed relatively short galactan backbone with a mixture of β-1,6 and β-1,3 links (Tan et al., 2004; Tan et al., 2010). The side chains were also short and contained a mixture of galactose, arabinose, glucuronic acid, and rhamnose. How well these structures reflect native structures is unclear. Larger glycans were found in radish roots, wheat flowers, and *Arabidopsis* leaves based on mass spectrometric analysis of enzymatically released AGP glycans (Haque et al., 2005; Tryfona et al., 2010; Tryfona et al., 2012). The evidence from these structural glycan studies and characterizations of GTs active on AGPs suggest a β-1,3 galactan backbone of varying length with β-1,6 side chains containing mainly galactan and arabinose residues with some additional sugar residues also found in pectin.

We hypothesize that hydrolytic enzymes acting on AGP glycans may also be involved in the synthesis of the arabinogalactan chains or their modification in the cell wall matrix. There is precedent for apoplastic post-deposition modification of xyloglucan by hydrolases (Gunl et al., 2011; Gunl and Pauly, 2011), but secretory pathway modification as known from N-glycan biosynthesis should also be considered. Based on the description of hydrolase enzymes in the carbohydrate-active enzyme database (CAZy), glycoside hydrolase family 43 (GH43) was identified as a potential group with AGP glycan hydrolyzing activity (Kotake et al., 2009; Geshi et al., 2013; Mewis et al., 2016). The GH43 family is conserved across prokaryotes and eukaryotes; it currently contains 15 620 GH43 enzymes, making it one of the largest known hydrolase families. The family is divided into α-L-arabinofuranosidase, β-D-xylosidase, α-L-arabinanase, and β-D-galactosidase groups (Mewis et al., 2016). Several GH43 enzymes from prokaryotes and eukaryotes have recently been characterized because of their potential in biomass degradation and other biotechnological applications (Jordan et al., 2013; McCleary et al., 2015). Based on their diverse identified enzymatic activities and amino acid motifs, the GH43s have been further subdivided into 37 subclades (Mewis et al., 2016). However, no GH43 family enzyme from plants has yet been characterized. The *Arabidopsis thaliana* genome contains two genes encoding amino acid sequences with similarity to GH43 enzymes belonging to the GH43 subfamily 24, named GH43A and GH43B (Mewis et al., 2016). Here, we characterize these GH43 enzymes and describe their role in cell expansion and root growth in *Arabidopsis thaliana*.

## Results

### *GH43* null mutants are defective in cell expansion

The *Arabidopsis thaliana* genome encodes two putative glycosyl hydrolase 43 family enzymes according to the latest version of the *Arabidopsis* Information Resource database (TAIR; https://www.arabidopsis.org/). Based on publicly available expression data, both genes are expressed throughout plant development with high expression in roots and stems. To investigate the functional role of these GH43 enzymes, we first obtained mutants carrying exon T-DNA insertions in *GH43A* and *GH43B* (Fig 1a). Initial analysis of the single *gh43* mutants and the double *gh43a-1/gh43b-1* mutant (*gh43null* henceforth) revealed no obvious visual phenotypes in seedlings or mature plants (Fig 1b and Supplemental Fig 1). Since *GH43* genes are expressed in seedlings and young roots according to publicly available expression data (eFP browser: http://bar.utoronto.ca/), we focused our subsequent phenotyping on seedlings grown on nutrient media agar plates.

**Figure 1.**
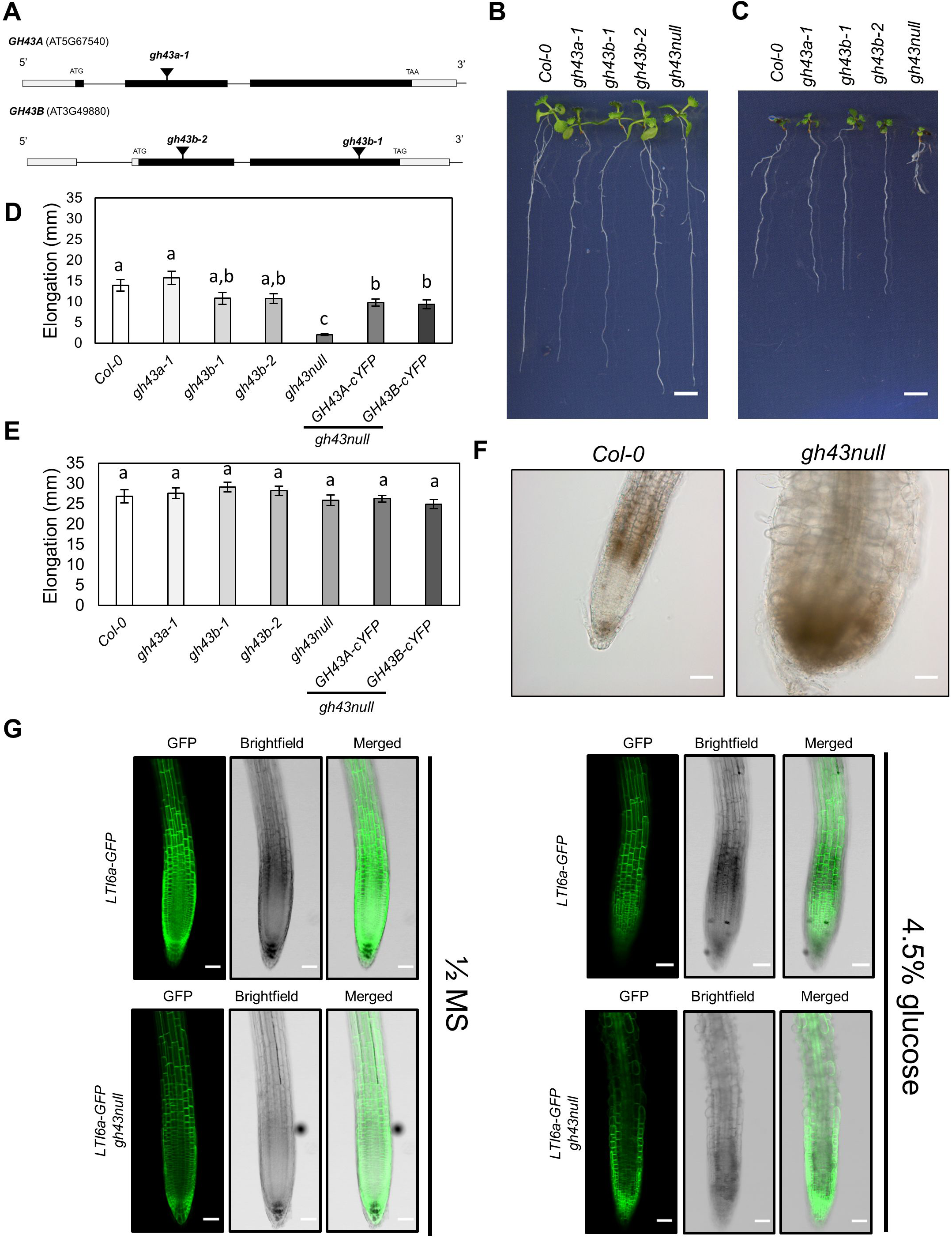
Characterisation of the *gh43* Arabidopsis T-DNA insertion lines. **(A)** Schematic diagram of the *GH43A* and *GH43B* gene structure and the T-DNA insertion sites. The white and black boxes indicate untranslated and translated regions, respectively. **(B)** Col-0 and *gh43* seedlings grown on nutrient media without sugar for 4 days and then moved to media without sugar for 6 days. Scale bar = 5mm **(C)** Col-0 and *gh43* seedlings grown on nutrient media without sugar for 4 days and then moved to media with 4.5% glucose for 6 days. Scale bar = 5mm **(D)** Root elongation of Col-0 and *gh43* seedlings grown on nutrient media without sugar for 4 days and then 6 days on media with 4.5% glucose. Values are means ±SE (*n* = 25 – 28 biological replicates). Means not sharing a common letter are significantly different at P < 0.05, as determined by Tukey’s test after one-way ANOVA. **(E)** Root elongation of Col-0 and *gh43* seedlings grown on nutrient media without sugar for 4 days and then 6 days on media without glucose. Values are means ±SE (*n* = 25 – 28 biological replicates). Means not sharing a common letter are significantly different at P < 0.05, as determined by Tukey’s test after one-way ANOVA. **(F)** Light microscopy images of Col-0 and *gh43null* root tip after seedlings were grown 4 days on nutrient media followed by 6 days on 4.5% glucose media. Scale bar = 100 µm. **(G)** Confocal laser scanning microscope images of Col-0 and *gh43null* roots stably expressing the *LTI6a-GFP* plasma membrane marker. Images were obtained 10 hours after moving 4-day old seedlings to nutrient media without (left) and with 4.5% glucose (right). Scale bars = 100 µm.

Several classic cell wall mutants in Arabidopsis show enhanced or conditional root growth defects on media containing 4.5 % exogeneous sugar (Benfey et al., 1993; Hauser et al., 1995). We therefore grew the *gh43* mutants on nutrient media without sugar for four days and then moved them to media containing 4.5 % glucose for six days. During the six days on glucose media, the *gh43a* and WT roots elongated at similar rates. However, root growth was slightly reduced in the *gh43b-1* and *gh43b-2* single mutants, and severely inhibited in *gh43null* (Fig 1c, and 1d). No change in growth rate was observed in controls moved to media without sugar (Fig 1e). On sugar-containing media, the *gh43null* root epidermal cells exhibited clear swelling and loss of anisotropic growth (Fig 1f). A time course experiment comparing WT and *gh43null* lines expressing the plasma membrane marker LTi6a-GFP revealed that the swelling was detectable in the cell elongation zone already after 6 hours and became obvious after 10 hours (Fig 1g and Supplemental Fig 2). This cell expansion defect is not observed on media containing 4.5 % sorbitol or 100 mM NaCl, showing that the phenotype cannot be caused by an osmotic effect alone or salt stress (Supplemental Fig 3). To confirm the causal gene defect responsible for the sugar-inducible loss of anisotropic growth, we performed complementation experiments by introducing either *GH43A-cYFP* or the *GH43B-cYFP* under the control of their native promoters into *gh43null* background. Both constructs rescued the root growth phenotype (Fig 1d).

### GH43A and GH43B are Golgi localized exo-β1,3-galactosidases

The *pGH43A:GH43A-cYFP* and *pGH43B:GH43B-cYFP* lines were used to investigate the subcellular location of the GH43s. In the expanding root cells of four-day old seedlings, both GH43A-cYFP and GH43B-cYFP appeared to localize to the Golgi apparatus (Fig 2a). To confirm the Golgi localisation, the lines were crossed with lines carrying the *cis-*Golgi marker SYP32-mCherry or the *trans-*Golgi network marker SYP43-mCherry (Fig 2a) (Uemura et al., 2004; Geldner et al., 2009). Both GH43 proteins co-localised with the Golgi markers. The overlap with SYP32-mCherry was higher than that with SYP43-mCherry, indicating a slight enrichment in the *cis-*Golgi (Fig 2b and 2c). In further support of the Golgi localisation, both Arabidopsis GH43s were identified in Golgi-enriched cell extracts in global proteomics experiments (Parsons et al., 2012; Parsons et al., 2019).

**Figure 2.**
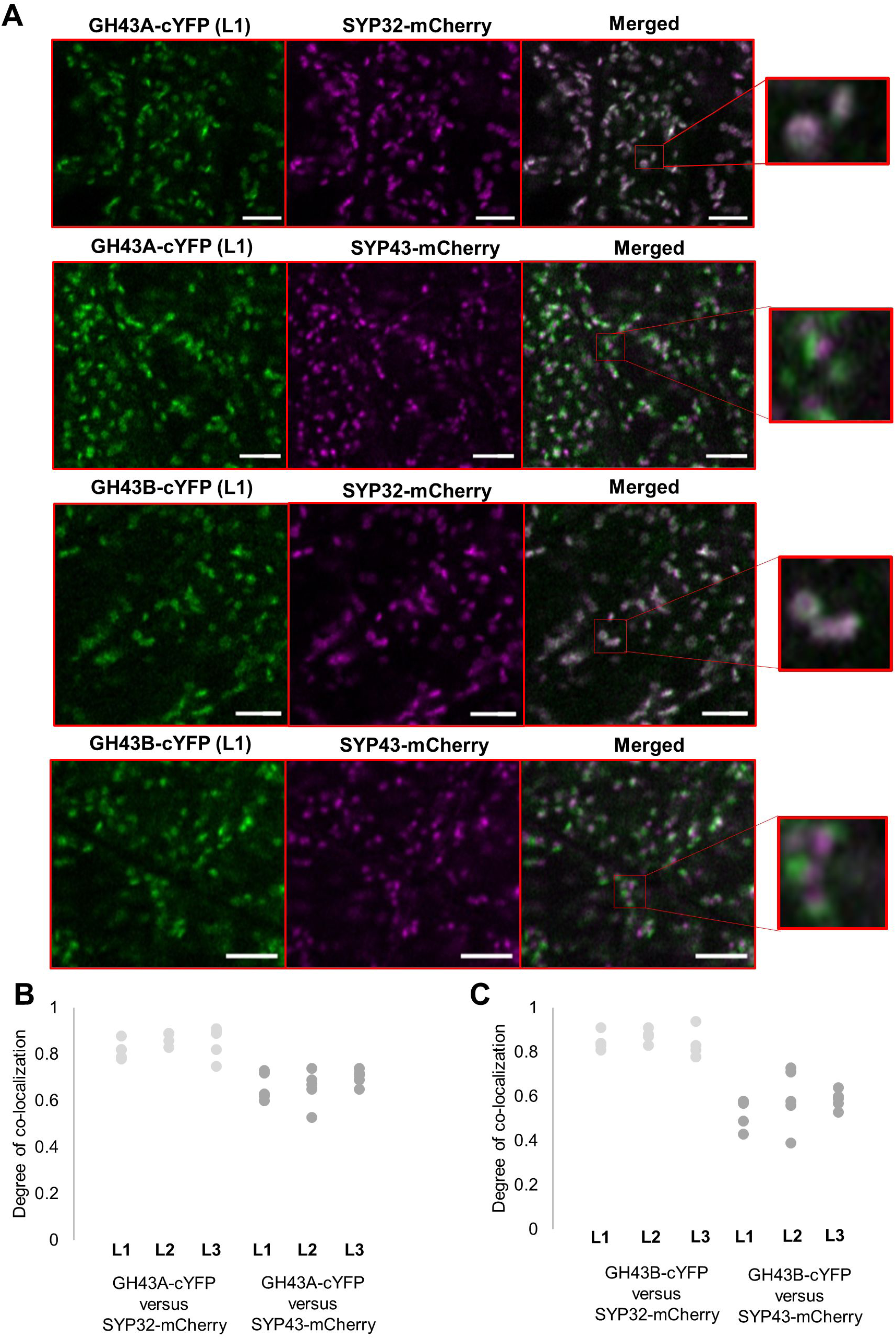
Co-localization of GH43A-YFP and GH43B-YFP with mCherry Golgi markers. **(A)** Confocal laser scanning microscope images of root epidermis cells in Arabidopsis seedlings stably expressing pGH43A::GH43A-cYFP or pGH43B::GH43B-cYFP and the cis-Golgi marker SYP32-mCherry or the Trans Golgi Network (TGN) marker SYP43-mCherry. Images were captured with a Zeiss LSM880 confocal microscope. Scale bars = 5µm. **(B)** Degree of co-localization of pGH43A::GH43A-cYFP with SYP32-mCherry or SYP43-mCherry in the root elongation zone epidermis. Images were captured from three independent GH43A-cYFP lines (L1-L3) and five seedlings per line. The values represent the degree of Pearson’s correlation between the YFP and RFP channel. **(C)** Degree of co-localization of pGH43B::GH43B-cYFP with SYP32-mCherry or SYP43-mCherry in the root elongation zone epidermis. Images were captured from three independent GH43A-cYFP lines (L1-L3) and five seedlings per line. The values represent the degree of Pearson’s correlation between the YFP and RFP channel.

The predicted hydrolase domains of the two Arabidopsis GH43 proteins exhibit over 90 % similarity at the amino acid sequence level, suggesting similar enzyme activity (Supplemental Fig 4). Sequence similarity also indicates that these proteins belong to the GH43 subfamily of enzymes with β-1,3-galactosidase activity (Ichinose et al., 2005; Ichinose et al., 2006; Kotake et al., 2009; Jiang et al., 2012; Mewis et al., 2016). To test for this activity, we expressed GH43A and GH43B in *Escherichia coli* and purified the recombinant proteins using nickel ion affinity chromatography (Supplemental Fig 5a). The activity of the recombinant GH43s was assayed against various of β-1,3-galactan di- and trisaccharides (Fig 3). Both GH43 proteins hydrolysed β-D-Galp-(1→3)-β-D-GalpOMe, but not β-D-Galp-(1→6)-β-D-GalpOMe. To confirm that the observed β-1,3-galactosidase activity was due to the GH43 enzymes, we generated GH43 enzymes with mutations in the predicted active site. The mutation sites were selected based on sequence similarity with the active site of a *Clostridium thermocellum* GH43 protein whose structure was solved by crystallography (Supplemental Fig 5b) (Jiang et al., 2012). Of the mutated versions, the GH43B^EQ224^ had minor activity towards β-D-Galp-(1→3)-β-D-GalpOMe, while GH43B^EQ224,338^ and GH43B^337-339DEL^ were inactive towards β-D-Galp-(1→3)-β-D-GalpOMe (Supplemental Fig 5c). These assays confirmed that the Arabidopsis GH43s are β-1,3-galactosidases and support the conservation of active sites between bacteria and plants.

**Figure 3.**
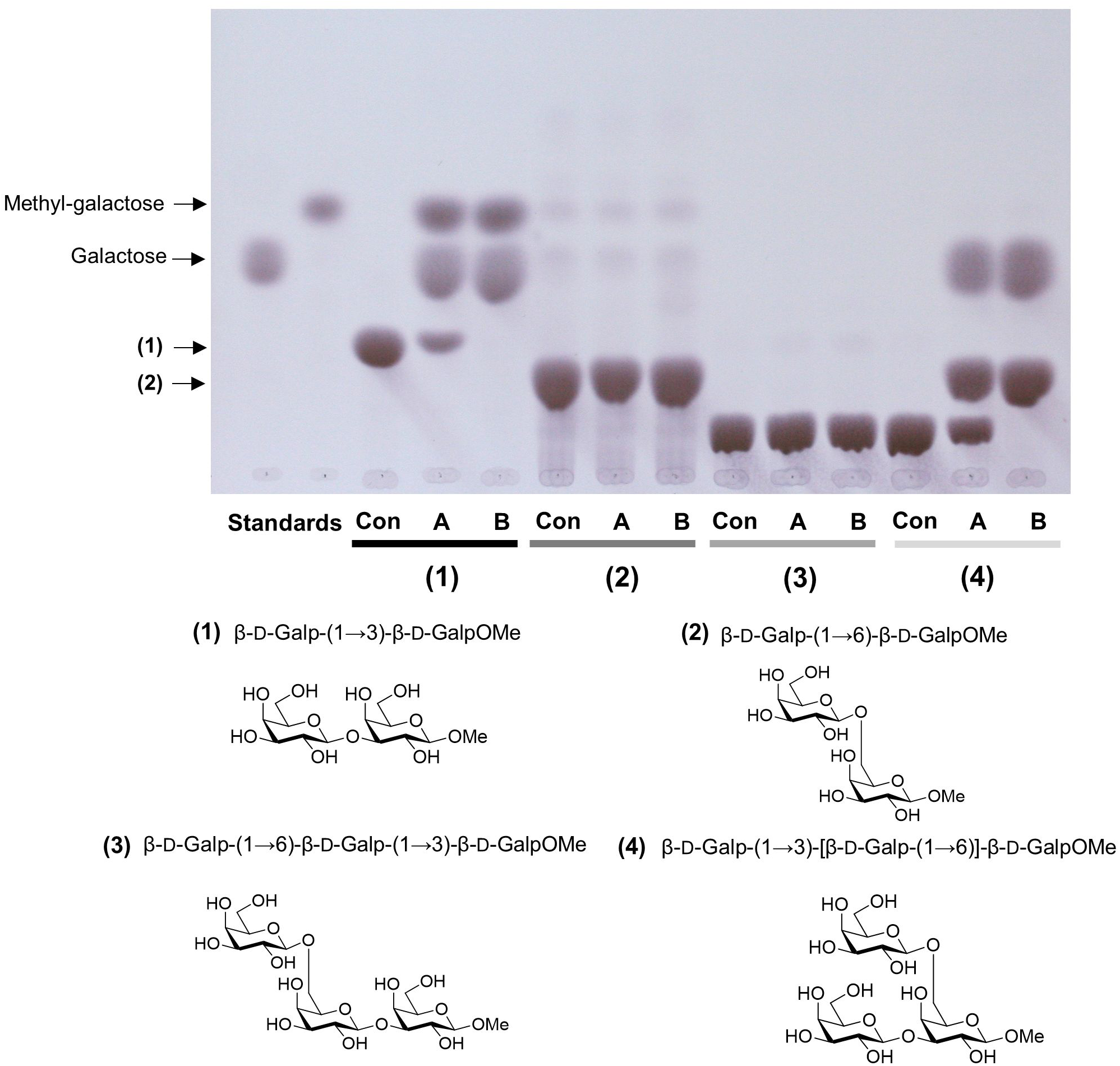
Thin layer chromatography (TLC) analysis of sugars released by recombinant Arabidopsis GH43 activity. Hydrolytic activity of heterologous GH43A and GH43B against **(1)** Methyl β-D-galactopyranosyl-(1→3)-β-D-galactopyranoside, **(2)** Methyl β-D-galactopyranosyl-(1→6)-β-D-galactopyranoside, **(3)** Methyl β-D-galactopyranosyl-(1→6)-β-D-galactopyranosyl-(1→3)-β-D-galactopyranoside and **(4)** Methyl β-D-galactopyranosyl-(1→3)-[β-D-galactopyranosyl-(1→6)]-β-D-galactopyranoside. The digestion products were separated with TLC. Con= substrate in buffer without enzyme, A = GH43A and B = GH43B.

β-1,3-galactan in plant cell walls is mainly found on AGPs (Du et al., 1996). Therefore, to investigate the presence of possible GH43 substrates in Arabidopsis cell walls, we sequentially extracted cell wall glycoproteins from Arabidopsis leaves using 0.2M CaCl_2_, 50 mM CDTA, 0.5M NaCO_3_, and 4M NaOH, and analysed the extracts by thin layer chromatography (TLC) before and after treatment with the GH43 proteins (Supplemental Fig 6a). Interestingly, the recombinant GH43s did not release galactose from any of the cell wall fractions, suggesting that the β-1,3-linkages were inaccessible. The recombinant GH43s also failed to release galactose from Gum Arabic, which is rich in AGPs (Supplemental Fig 6b) (Akiyama et al., 1984). Even partial deglycosylation of the AGPs was insufficient to allow the GH43 proteins access to the β-1,3-linkages (Supplemental Fig 6b). These observations suggested that the β-1,3-linkages of mature AGPs may be protected against GH43 activity.

To investigate the enzymatic mechanism of the Arabidopsis GH43 proteins in more depth, we synthesized β-1,3 galactan oligosaccharides with a β-1,6-galactose branch on the reducing and non-reducing ends of β-D-Galp-(1→3)-β-D-GalpOMe. The GH43s were unable to hydrolyze the β-D-Galp-(1→3)-β-D-GalpOMe when the non-reducing end bore a β-1,6-galactose unit, but did hydrolyze it when the substitution was at the reducing end (Fig 3). Steric hindrance due to the presence of side chains at the non-reducing end of the β-1,3 galactan thus protects against hydrolysis catalysed by GH43 proteins. We also tested the ability of GH43s to hydrolyse β-1,3- or β-1,4-glucans, but observed no activity towards these polymers (Supplemental Fig 6c). These localisation and recombinant enzyme assays established the two Arabidopsis GH43s as Golgi exo-β-1,3 galactosidases with quite strict substrate specificity.

### *gh43* null mutants have altered cell wall structure

WT and *gh43null* seedlings had similar contents of cellulose, hemicellulosic and pectic sugars (Supplemental Fig 7, Supplemental Table 1). The *gh43null* root swelling thus cannot be explained by changes in the levels of the main cell wall polymers, although it is possible that more localized cell-type specific defects were masked by the analysis of total tissue extracts. Based on the GH43 proteins’ Golgi localisation, exo-β1,3-galactosidase activity and inactivity towards AGP-containing plant extracts, we hypothesised that the they are involved in the synthesis of the arabinogalactans in the Golgi. It is plausible that their native targets occur only found in the Golgi and represent a small fraction of the total β-1,3-galactan pool. This hypothesis is consistent with the inactivity of the GH43 proteins towards β-1,3 galactans substituted with a β-1,6 galactan at the non-reducing end, and suggests that GH43s may be involved in the processing of the AGPs during biosynthesis.

We hypothesised that defects in arabinogalactan synthesis in the *gh43null* background would cause changes in the structure and/or extractability of AGPs. To investigate this possibility, we first quantified the levels of cell wall-bound AGPs (isolated by removing the CaCl_2_-soluble fraction followed by AIR1 treatment) and AGPs soluble in 0.18 M CaCl_2_ in WT and *gh43null* seedlings using β-Yariv. Yariv phenyl glycosides selectively bind to the β-1,3 galactans of AGPs, enabling their spectrophotometric quantification (Kitazawa et al., 2013). The fraction of cell wall-bound AGPs in *gh43null* was higher than in the WT (Fig 4a), where as a tendency for the opposite was observed for the soluble fraction (Fig 4b). We thus hypothesized that structural differences in the AGP glycans of the *gh43null* mutant may cause some AGPs to bind more tightly to the cell wall.

**Figure 4.**
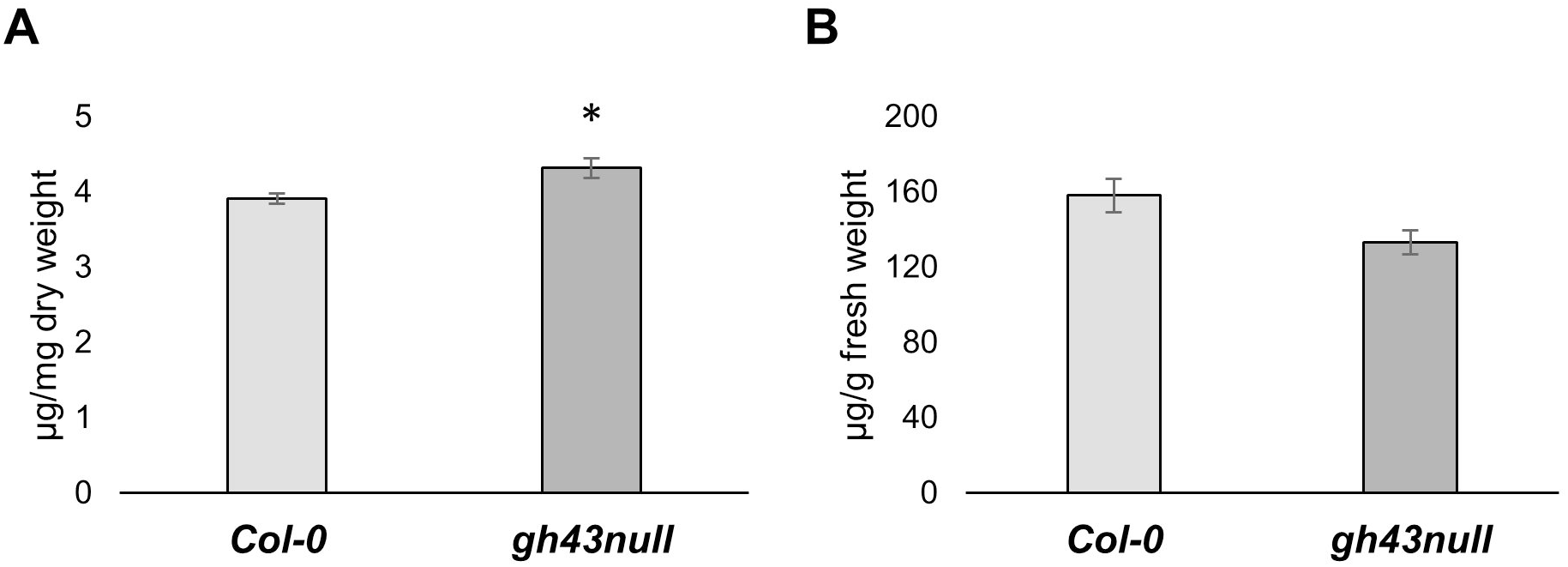
Spectrophotometric quantification of cell wall bound and soluble arabinogalactan proteins with β-Yariv. **(A)** β-Yariv binding to wild type (Col-0) and *gh43null* cell walls from seven-day old seedlings after removal of CaCl_2_ and AIR1 soluble fractions. Amount of cell wall bound β-Yariv was quantified against a Gum Arabic standard. Values are means ±SE. *= P < 0.05 (unpaired *t* test, *n* = 5 biological replicate pools of seedlings). **(B)** β-Yariv binding to the CaCl_2_ soluble fraction from wild type (Col-0) and *gh43null* seven-day old seedlings. Amount of bound β-Yariv was quantified against a Gum Arabic standard. Values are means ±SE. (*n* = 5 biological replicate pools of seedlings).

AGPs bound to cell walls may affect matrix properties and conceivably also cell expansion. To test whether this could explain the root swelling phenotype, we sequentially extracted cell wall fractions from 7-day old WT and *gh43null* seedlings using 180 mM CaCl_2_, 50 mM CDTA, and 4 M NaOH/26.5 mM NaBH_4_. Equal amount of each fraction was analysed on a Comprehensive Microarray Polymer Profiling (CoMPP) microarray designed to determine different sugar epitopes in complex plant extracts (Moller et al., 2012). Similar polymer profiling has also previously been employed to demonstrate cell wall matrix separability features (Harholt et al., 2012; Tan et al., 2013). The CoMPP microarray contained monoclonal antibodies specific for pectin, xylan, xyloglucan, mannan, crystalline cellulose, extensin and AGP epitopes (Supplemental Table 2). For epitopes released with 180mM CaCl_2_, the JIM13, LM20 and RU2 antibody signals of *gh43null* were significantly weaker than those of the WT (Table I). Pectin antibodies LM20 and RU2 bind partially methylesterified homogalacturonan and the rhamnogalacturonan-I (RG-I) backbone, respectively (Verhertbruggen et al., 2009; Ralet et al., 2010), while JIM13 recognises an AGP epitope (Yates et al., 1996). For epitopes released with 50 mM CDTA, the LM5, LM15 and LM18 signals of the *gh43null* were significantly stronger than those of the WT (Table I). Pectin epitope antibody LM5 binds to β1,4-galactan, LM18 to partially methylesterified homogalacturonan, and LM15 to xyloglucan (Jones et al., 1997; Marcus et al., 2008; Verhertbruggen et al., 2009). The cell wall fraction exhibiting the most pronounced differences between *gh43null* and WT was the extracted with 4 M NaOH/26.5 mM NaBH_4_ released epitopes (Table I). In accordance with the β-Yariv assay results (Fig 4a), the signals from the AGP antibodies JIM13, LM2, LM14 and MAC207 were all significantly more intense in *gh43null* (Table I) (Smallwood et al., 1996; Yates et al., 1996; Ruprecht et al., 2017). An increased signal intensity in *gh43null* was also observed for the pectin (galactan) antibody LM5, the xyloglucan binding LM25, and the CBM3a crystalline cellulose and JIM20 extensin antibodies (Smallwood et al., 1994; Verhertbruggen et al., 2009; Hernandez-Gomez et al., 2015). These CoMPP results established that the defect caused by the loss of GH43 β-1,3-galactosidase activity causes changes in the abundance of cell wall-associated AGPs.

**Table I.**
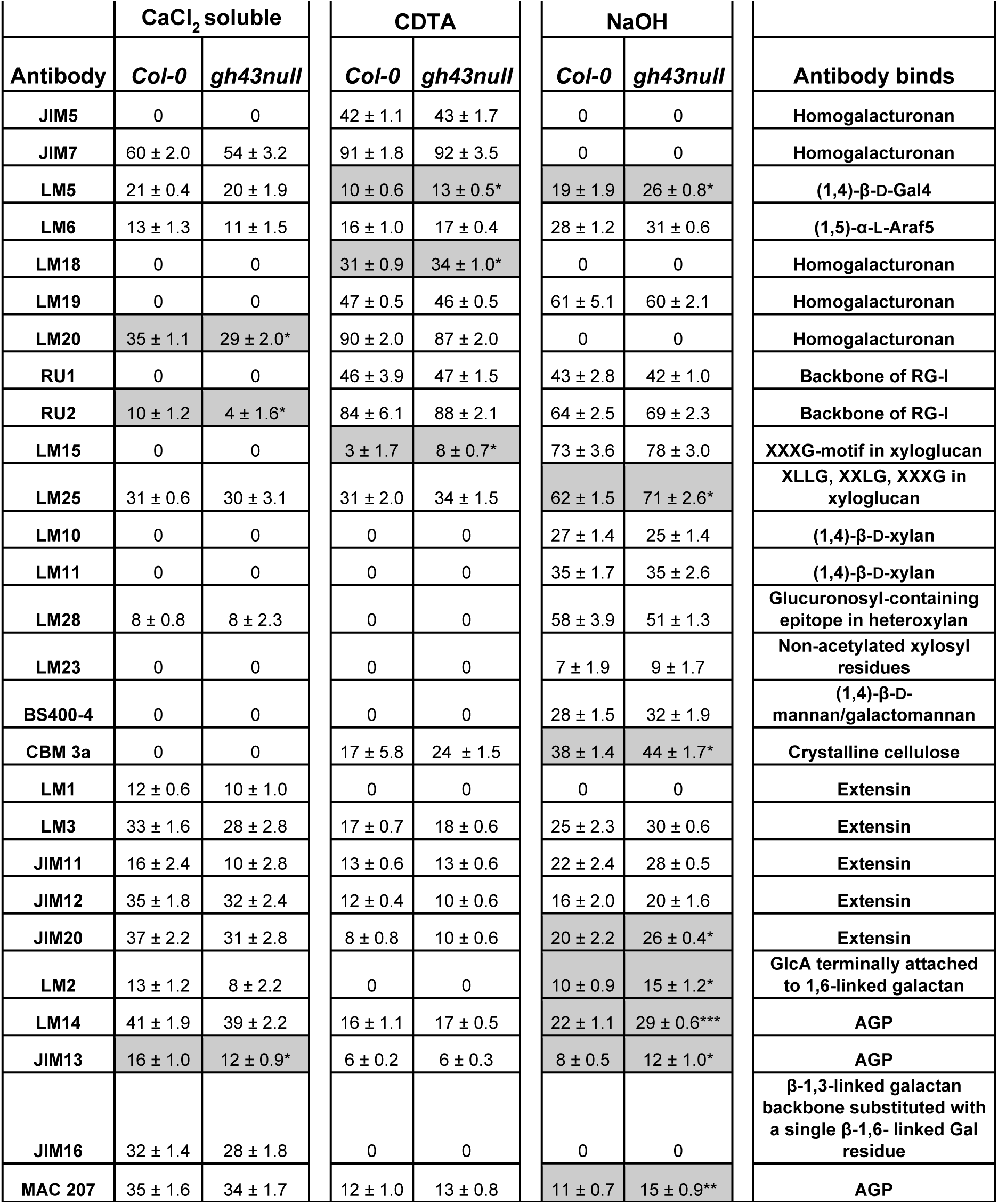
Comprehensive Microarray Polymer Profiling (CoMMP) of sequentially extracted cell walls from seven-day old Col-0 and *gh43null* seedlings. Cell wall fractions extracted with 180mM CaCl_2_, 50 mM cyclohexanediamine tetraacetic acid (CDTA) and 4M NaOH/26.5mM NaBH_4_. The values represent the signal intensity of the fluorescent secondary antibody. Values are means ±SE. Grey cells mark significant differences *= P < 0.05, **= P <0.01, ***= P <0.001 (Student’s *t* test, *n* = 5 biological replicates with pooled seedlings per replicate).

## Discussion

β1,3-galactan in plant cell walls is found primarily on AGPs (Du et al., 1996). We identified and characterized the enzymatic activity of two Golgi-localized GH43 exo-β1,3-galactosidases and discovered that their activity influences the levels of cell wall associated AGPs. Functional characterisation of plant GH43 enzymes has not previously been reported. The active site of the prokaryotic GH43 from *Clostridium thermocellum* and the Arabidopsis GH43s is conserved (Supplemental Fig 5). Despite this catalytic similarity, the Arabidopsis GH43s are involved in cell wall biosynthesis whereas their prokaryotic and fungal counterparts are involved in plant cell wall degradation (Ichinose et al., 2005; Ichinose et al., 2006; Kotake et al., 2009; Jiang et al., 2012; Mewis et al., 2016). The functional role of GH43 activity during root growth was revealed by experiments using nutrient media supplemented with 4.5% glucose. On such media, root growth is modestly reduced in single *gh43b* mutants and dramatically reduced in the *gh43null* double mutant (Fig 1). Nutrient media with 4.5% sugar was also used in a classical genetic screen for conditional root growth mutants of Arabidopsis (Benfey et al. 1993; Hauser et al. 1995). Several mutants identified in this way were subsequently shown to be defective in cell wall biosynthesis genes including the glycosylphosphatidylinositol-anchored plasma membrane protein *COBRA* (Schindelman et al., 2001), the chitinase-like protein *CTL1* (Sanchez-Rodriguez et al., 2012) and the cellulose synthase interacting 1 (*CSI1*) (Gu et al., 2010). The mechanism responsible for the sugar-induced root growth defects in these cell wall mutants and the *gh43* mutants remains unclear, but may involve cell wall acidification of sugar-activated proton pumps and/or disturbance of hormone homeostasis (Niittyla et al., 2007; Ljung et al., 2015; Yeats et al., 2016). The root cell expansion defects of *gh43null* become noticeable in the elongation zone within 6 hours of transferring seedlings to sugar media (Supplementary Fig 2). Quantification of cell wall-associated AGPs with β-Yariv (Fig 4), and the CoMPP analysis of cell wall fractions from untreated seedlings (Table I) confirmed that the *gh43null* cell walls differ from WT even before the sugar treatment. Thus, a pre-existing cell wall defect is likely to underlie the sugar induced swelling.

Investigation of the *gh43null* cell wall defect revealed a connection to AGPs. We observed AGP epitopes in all sequentially extracted cell wall fractions (Table I). The increase in the levels of AGPs extracted from the *gh43null* with 4 M NaOH (i.e. AGPs not released by 180mM CaCl_2_ or 50mM CDTA) suggests that GH43s play a role in adjusting the cell wall affinity of AGPs. The CoMPP microarray analysis of cell wall fractions also revealed subtle differences in the abundance of pectin and xyloglucan epitopes in the *gh43null* mutant (Table I). We hypothesise that these changes reflect modifications of the cell wall matrix resulting from initial changes in the AGPs. Based on the inactivity of GH43s towards β-1,3 galactans with β-1,6 branching (Fig 3), and the lack of galactose release from extracted cell wall glycoproteins or AGP-rich gum arabic (Supplemental Fig 6), we propose that the Arabidopsis GH43s are required for β-1,3-galactan processing during AGP maturation in the Golgi. It is conceivable that GH43s regulate the length of the β-1,3-galactan backbone (and thus the number of side chains that can attach to it), and/or modify the backbone length to make it accessible to other enzymes. The Golgi-associated α-mannosidase I, which is involved in structuring N-glycosylation, has been demonstrated to perform proteoglycan trimming of this sort in Arabidopsis (Liebminger et al., 2009). A similar AGP glycan structuring role could be envisaged for GH43s.

Accumulating evidence from several plant species supports the existence of cell wall matrix-bound AGPs, primarily associated with pectin (summarised by Tan et al. (2013)). Pectin can be subdivided into homogalacturonan (HG), xylogalacturonan (XGA), apiogalacturonan, rhamnogalacturonan type I (RG-I), and rhamnogalacturonan type II (RG-II) (Harholt et al., 2010). Of these, HG and RG-I are the most abundant in Arabidopsis primary walls, accounting for ~65% and 20-35% of the total pectin content, respectively (Mohnen, 2008). HG is a linear α-1,4-linked galacturonic acid that is partially methylesterified and can be O-acetylated. RG-I consists of a α-1,4-D-galacturonic acid-α-1,2-L-rhamnose backbone with several side chains (Harholt et al., 2010). The RG-I side chains include galactans and arabinans as well as arabinogalactan type I (AG-I), and arabinogalactan type II (AG-II) polysaccharide chains (Yapo, 2011). The AG-II structure resembles that of AGP glycans and it may be that some of the RG-I attached oligosaccharides actually derive from AGP glycans. In Arabidopsis, the AGP ARABINOXYLAN PECTIN ARABINOGALACTAN PROTEIN1 (APAP1) binds covalently to RG-I/HG and was shown to increase the extractability of pectin and hemicellulosic immunoreactive epitopes, suggesting that it acts as a structural cross-linker in plant cell walls for this AGP (Tan et al., 2013). Whether the covalent bond between APAP1 and RG-I is a result of transglycosylation taking place in the apoplast or synthesises as such in the Golgi is unknown. Interestingly, the CoMPP microarray results for the *gh43null* mutant revealed decreased homogalacturonan (LM20) and RGI-backbone (RU2) signals in the CaCl_2_ soluble fraction, and increased β-1,4-galactan (LM5) and homogalacturonan (LM18) signal in the CDTA fraction and β-1,4-galactan (LM5) in the NaOH fraction (Table I). Together, these results show that *gh43null* exhibits altered pectin extractability compared with the WT, and suggest that APAP1 and/or similar cell wall-bound AGPs could be targets of GH43 activity.

Pectin in expanding cells has been proposed to function as a mechanical tether between cellulose microfibrils (Höfte et al., 2012), and as a lubricant of microfibril movement during cell expansion (Cosgrove, 2014). Experimental support for direct interaction between cellulose and pectin in primary cell walls of Arabidopsis was provided by the comparison of never dried, dehydrated and rehydrated cell walls using multidimensional solid state nuclear magnetic resonance (NMR) spectroscopy (Wang et al. (2015). The NMR spectra showed cross peaks between cellulose and pectin indicating that some of the pectic sugars come into subnanometer contact with cellulose microfibrils. Based on these pectin models, AGP and pectin interactions would be expected to influence primary cell wall extensibility by restricting cellulose microfibril movement and/or cell wall matrix extension. This concept was first introduced in an early cell wall model proposed in the 1970s, which depicted covalent connections between pectin and structural proteins (Keegstra et al., 1973). However, strong evidence of the role of structural cell wall proteins has been slow to emerge and therefore many current cell wall models omit them. The involvement of GH43s in the modification of AGP association with cell walls breathes new life into the concept of structural cell wall proteins, and establishes a previously unknown β-1,3-galactan trimming process during primary plant cell biosynthesis and cell growth. Cell wall bound AGPs may play an important role as structural matrix proteins preventing cellulose microfibril coalescence, simultaneously hindering other cell wall polymers from binding to cellulose. In this model, more tightly bound AGPs could impair wall stiffening during cell expansion, and thus contribute to the *gh43null* root swelling phenotype. This interpretation would imply that the distribution, measured as extractability, of other polymers in the wall would also change and this seems indeed to be the case in the *gh43null* mutant.

## Methods

### Plant material and growth conditions

*gh43a-1* (GABI_062D83), *gh43b-1* (Salk_002830), *gh43b-2* (SALK_087519) were ordered from NASC and genotyped (Supplemental table 3). The T-DNA insert locations were determined by sequencing (Fig 1a). Mutant and wild type plants were grown on soil at 22 °C with a photoperiod of 16 h light and 8 h dark at 65% relative humidity.

### Plant vector construction and transformation

The genomic sequences of GH43A and B were amplified using the primers in Supplemental Table 3. The amplified pieces were cloned in the pDONR207 plasmid via Gibson assembly and recombined into pHGY (Kubo et al., 2005). Both constructs were transformed into *gh43null.* Stably transgenic plants were generated by *Agrobacterium tumefaciens*–mediated transformation. Transgenic seedlings were selected based on hypocotyl elongation on 0.9% agar ½ MS plates supplemented with 30 μg/mL hygromycin. Homozygous lines were generated based on hygromycin segregation and detection of the YFP signal by confocal microscopy.

### Root elongation/complementation experiments

Seeds were surface sterilized with 70% ethanol and grown on ½ MS agar plates (0.9% agar, 5 mM MES, pH 5.7) for 4 days. After this period, the seedlings were moved to ½ MS plates, ½ MS plates containing either 4.5% glucose or 4.5% sorbitol or 100 mM NaCl plates. The seedlings were then allowed to grow for 6 days. The difference in root elongation from day 0 on the new plate to day 6 was quantified using Image J (https://imagej.nih.gov/ij/).

### GH43 protein localization and root imaging in the LTi6a-GFP background

The lines *pGH43A*::*GH43A-cYFP* and *pGH43B*::*GH43B-cYFP* co-expressing the marker lines SYP32-mCherry or SYP43-mCherry were generated by plant crossing (Uemura et al., 2004; Geldner et al., 2009). For observation, seedlings were grown vertically on plates containing ½ MS media with 0.9% plant agar for 5 days and then observed under a Zeiss LSM880 confocal laser scanning microscope. A Zeiss LSM880 confocal laser scanning microscope with an Airyscan detector and a LD LCI Plan-Apochromat 40x/1.2 Imm AutoCorr DIC M27 water immersion objective was used.

The *LTI6a-GFP* line was crossed in the *gh43null* mutant background and homozygous lines were generated (Grebe et al., 2003). The seedlings were grown for 4 days on ½ MS plates and then moved to ½ MS plates or ½ MS plates with 4.5% glucose. The roots were imaged with a Zeiss LSM880 confocal microscope.

### Heterologous expression and purification of the GH43 proteins

The *Arabidopsis* GH43 coding sequences were amplified from 121-1401 bp (AT5G67540.1, GH43A) and 115-1401 bp (AT3G49880, GH43B) to avoid the predicted membrane domain. The proteins were expressed in Rosetta^tm^ (DE3) *E.coli* cells using a pET24d His10SUMO vector. The crude protein extract was purified by Ni-NTA, then the His10SUMO tag was cleaved and the GH43 protein isolated by Ni-NTA chromatography (Supplemental Fig 5a).

### Synthesis of methyl β-D-Galactopyranoside substrates 1-4 (Fig 3)

Chemical synthesis of methyl 3-O-β-D-galactopyranosyl-β-D-galactopyranoside (1) (Kovac et al., 1985), methyl 6-O-β-D-galactopyranosyl-β-D-galactopyranoside (2) (Kovac et al., 1984), and methyl 3,6-di-O-(β-D-galactopyranosyl)-β-D-galactopyranoside (4) (Kaji et al., 2010) have been reported in literature. However, methyl β-D-galactopyranosyl-(1→6)-β-D-galactopyranosyl-(1→3)-β-D-galactopyranoside (3) has not previously been reported. The strategy for chemical synthesis of the substrates 1-4 is outlined in Supplemental Fig 8

### GH43 protein activity assays

The enzymes’ buffer was changed to 10 mM MOPS (pH 7) using the PD Spintrap™ G-25 (GE Healthcare, 28-9180-04) protocol. To determine the activity towards the β-galactan substrates (Fig 4) and β-glucan substrates (Supplemental Fig 6C), 1 µg GH43A and 4 µg GH43B were incubated with 100 µg of substrate (15 µl total volume), overnight at 30 °C while shaking. For tests using the inactive enzymes, 1 µg was loaded (Supplemental Fig 5c). Gum Arabic (200 mg/ml) was hydrolysed in 0.1 M TFA hydrolysed at 80 °C for 45 minutes (loading 10 µl on a TLC plate). The TFA was then removed using a speed vac by reducing the reaction mixture’s volume to around half of its original value. 30 mg of the resulting digestion product was further hydrolysed with an enzyme cocktail containing 1µl β-galactanase (E-BGLAN), 0.5µl β-glucuronosidase (E-BGLAEC), 18µl α-Fucosidase (E-FUCTM) and 20µl α-Arabinofuranosidase (E-ABFAN) from Megazyme in a 50 mM sodium acetate buffer pH5 at 37 °C for 4 hours (again, 10 µl was loaded on a TLC plate). The enzymes were then inactivated at 95 °C for 15 minutes, after which their buffer was changed to 10 mM MOPS (pH 7) using a PD MidiTrap G-10 column to 10mM MOPS pH7 to remove the sodium acetate and digestion products. The total sample was incubated with 2.5 µg GH43A or 10 µg of GH43B overnight while shaking at 30°C, then concentrated with a speed vac to approximately 10 µl and loaded onto a TLC plate.

Sequentially extracted cell wall material (10 mg/100 µl MOPS pH 7) was incubated with 2 µg GH43A or 7.5 µg of GH43B at 30 °C overnight while shaking. The samples were then concentrated with a speed vac to approximately 10 µl and loaded onto a TLC plate.

The products were loaded on a TLC Silica Gel 60 F254 TLC plate membrane and developed with 4:1:1 (1-Butanol:Acetetic acid: H_2_O). The membranes were visualized with 5 % Sulphuric acid and 0.5 % Thymol in 96 % ethanol at 105°C.

### AGP quantification with β-Yariv

Soluble AGP purification and quantification was performed according to Lamport (2013). For quantification of cell wall-associated AGPs, the cell wall pellet left after the removing CaCl_2_-soluble AGPs was flash frozen in liquid nitrogen and then lyophilized. The resulting material was ball milled, AIR1-treated (see below) and lyophilized. The lyophilized material was then incubated with β-Yariv for 4 hours at room temperature, and the β-Yariv absorbance was spectrophotometrically quantified according to Lamport (2013).

### Sequential extraction and Comprehensive Microarray Polymer Profiling (CoMPP)

Seeds were surface sterilized with 70 % ethanol and grown on ½ MS agar plates (0.9% agar, 5 mM MES, pH 5.7) for 7 days. The seedlings were then harvested, flash frozen, and ground in liquid nitrogen with a mortar and pestle. CaCl_2_-soluble glycoproteins were extracted with 0.18 M CaCl_2_ (2ml/g fresh weight) for 2 hours at RT and spun down at 4000gx10min. The pellet containing the cell wall material was freeze dried, ball milled, and AIR1-treated (see below) before further processing. The supernatant was precipitated with 4 volumes of ethanol at 4 °C for 16 hours and spun down at 2000g*2min. The pellet was further solubilized in 45 mM CaCl_2_ (2 ml/g fresh weight from starting material) and freeze dried. For the carbohydrate microarray, the soluble glycoproteins and cell wall material were further treated and analysed according to Moller et al. (2012). For determination of the monosugar composition, the fractions extracted with 50 mM CDTA and 4M NaOH/ 26.5mM NaBH_4_ fractions were dialysed against demineralised water using Visking® 12-14 kD dialysis tubing (45 mm).

### Wet chemical analysis of cell walls

The 7-day old seedlings were harvested, flash frozen and lyophilized. The dried material was ball milled, and the alcohol-insoluble residue (AIR1) was removed by incubating the material first for 30 minutes in 80% ethanol, and then for 30 minutes in 70 % ethanol at 95 °C, and finally in chloroform:methanol (1: 1) for 5 min at room temperature before washing with acetone. This sequence of incubations and washes constitutes the AIR1 treatment mentioned above. Starch (AIR2) was then removed according to Rende et al. (2016). The mono sugar composition was measured according to Latha Gandla et al. (2015). The crystalline cellulose content was quantified according to the Updegraff (1969).

## Supplemental data

**Supplementary Table 1.** Hemicellulosic/pectin sugar composition of sequential extracted cell wall fractions of 7-day old Col-0 and *gh43null* seedlings.

**Supplementary Table 2.** Binding specificity of the CoMPP antibodies

**Supplementary Table 3.** Primers used for genotyping the *gh43* lines and cloning of the *GH43-cYFP* constructs.

**Supplementary Figure S1.** Picture of ten-week-old *Col-0* and *gh43* lines.

**Supplementary Figure 2.** Confocal laser scanning microscope images of Col-0 and *gh43null* roots stably expressing the LTI6a-GFP plasma membrane marker.

**Supplementary Figure 3.** Col-0 and the *gh43* grown on 4.5% sorbitol or 100mM NaCl.

**Supplementary Figure 4.** Amino acid sequence alignment of GH43A and GH43B.

**Supplementary Figure 5.** Activity of heterologous expressed GH43B proteins with mutations in predicted active site.

**Supplementary Figure 6.** Activity of the heterologous expressed GH43 proteins towards Gum Arabic, partially digested Gum Arabic, sequentially extracted cell wall material and β1,3 or β1,4 glucan substrates.

**Supplementary Figure 7.** Crystalline cellulose content in 7-day old Col-0 and the *gh43null* seedlings.

**Supplementary Figure 8.** Chemical synthesis of the methyl β-galactopyronaosides.

## Acknowledgements

We would like to thank Mikael Lindberg from the Umeå University Protein Expertise Platform (PEP), for help with expressing the recombinant Arabidopsis GH43s. We would like to thank Dr. Junko Takahashi-Schmidt and the UPSC Biopolymer Analytical Platform for help with the cell wall monosugar analysis. This work was supported by The Swedish Foundation for Strategic Research (Value Tree), Bio4Energy (Swedish Programme for Renewable Energy), the UPSC Centre for Forest Biotechnology funded by VINNOVA and the Swedish Research Council for Sustainable Development (Formas).

## Author Contribution

P.N., B.P., M.S.M., B.J. and P.U. planned and performed experiments and analyzed data. T.N. planned experiments and analyzed data. P.N. and T.N. wrote the manuscript with help from the other authors.

## Parsed Citations

We would like to thank Mikael Lindberg fromthe Umeå University Protein Expertise Platform(PEP), for help with expressing the recombinant Arabidopsis GH43s. We would like to thank Dr. Junko Takahashi-Schmidt and the UPSC Biopolymer Analytical Platformfor help with the cell wall monosugar analysis. This work was supported by The Swedish Foundation for Strategic Research (Value Tree), Bio4Energy(Swedish Programme for Renewable Energy), the UPSC Centre for Forest Biotechnologyfunded by VINNOVAand the Swedish Research Council for Sustainable Development (Formas).

